# Multi-modal Monte Carlo MRI simulator of tissue microstructure

**DOI:** 10.1101/2025.04.03.646577

**Authors:** Michiel Cottaar, Zhiyu Zheng, Karla Miller, Benjamin C. Tendler, Saad Jbabdi

**Affiliations:** Wellcome Centre for Integrative Neuroimaging (WIN), Centre for Functional Magnetic Resonance Imaging of the Brain (FMRIB), University of Oxford, UK

## Abstract

Monte Carlo simulation of MRI signals is a powerful tool for modelling and quantifying tissue microstructure. While these methods have been used in MRI for decades, they typically focus on one aspect of microstructure and one type of MRI contrast at a time. In this paper, we present a new Monte Carlo MRI (MCMR) simulator for investigating the impact of tissue microstructure on arbitrary MRI sequences. A key feature of the proposed simulator is that substrates can incorporate multiple microstructural features (diffusion, T1/T2 relaxation, membrane permeability, local off-resonance fields, surface relaxation and magnetisation transfer) simultaneously. This provides a single forward model from tissue microstructure to MRI signals that captures a broad range of contrasts typically considered to be of different MRI “modalities”. We validate the results of the simulator by reproducing previous findings in well-established sequences, namely diffusion-weighted MRI, magnetisation transfer imaging, and gradient and spin echo sequences. The simulator is fully open source and packaged along with detailed documentation and tutorials.

## 1. Introduction

Magnetic Resonance Imaging (MRI) is a very flexible tool to image biological tissue. A wide variety of MRI modalities have been developed to probe different properties of the tissue microstructure (e.g., myelin density or cell size). However, combining information across modalities is non-trivial. Each MRI modality has a different contrast-forming mechanism and, hence, tends to come with its own microstructural model.

While all of these models aim to represent the same underlying tissue, they are often inconsistent with each other. For example, the binary spin-bath model commonly adopted when interpreting magnetisation transfer data (Henkelman *et al*., 1993; Manning, MacKay and Michal, 2021) assumes a single free water compartment that is perfectly mixed. This directly contradicts the assumption of non-exchanging, distinct pools of free water commonly adopted in diffusion MRI (Zhang *et al*., 2012; Novikov, Kiselev and Jespersen, 2018). However, exchange between such compartments is unlikely to be completely absent at the timescales probed by diffusion MRI, while at the same time fully completed at the timescale probed by magnetisation transfer.

Such simplifying assumptions are a necessity to derive quantitative microstructural parameters from the limited information available from a single MRI modality. These assumptions also allow for analytical solutions, which simplify fitting and generate insight on how the MRI signal depends on microstructural features. However, with the rise of more computational power as well as AI-based fitting tools (Jallais and Palombo, 2024), there is less of a need to limit ourselves to simplified microstructural models with analytical solutions. A more coherent analysis of multi-modal data would benefit from a single model that can jointly describe these multiple modalities.

With this aim in mind, we present here a new Monte Carlo MRI simulator that simultaneously captures multiple features of tissue microstructure that impact MRI measurements (Figure 1). These are:

**Figure 1.**
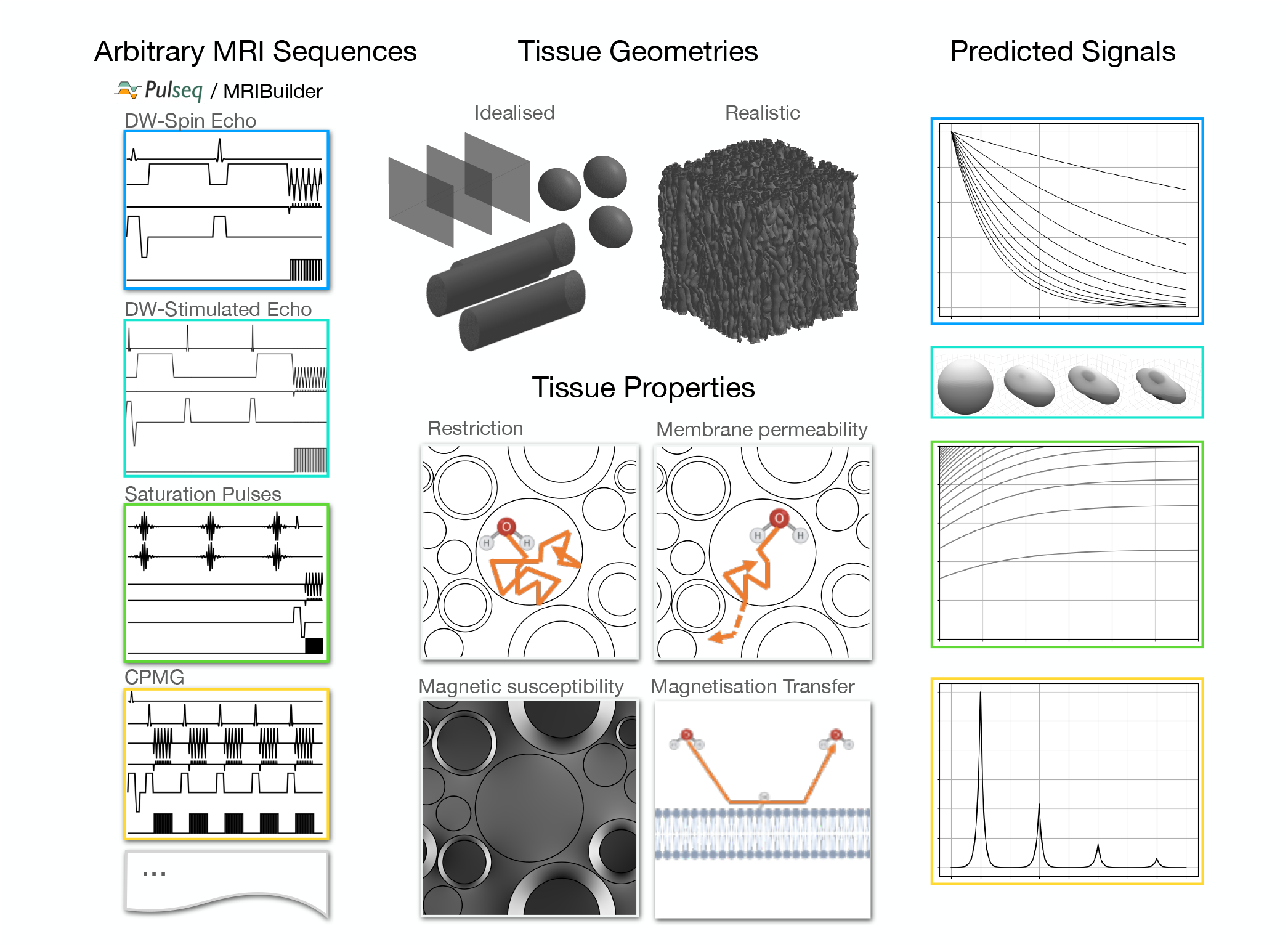
The MCMR simulator takes as input one or more MRI sequences (supplied as Pulseq files or the provided sequence builder software), a representation of the tissue microstructure geometry, and parameters describing T1/T2 relaxation, diffusion, membrane permeability, magnetic susceptibility, surface relaxation, and magnetisation transfer (Table 1). Given these inputs the MRI signals are predicted using Monte Carlo simulations.

1. Restriction of the random diffusion of water by tissue membranes (including permeability)
2. Alteration of the local magnetic field due to variations in magnetic susceptibility (e.g. myelin and iron in tissue)
3. Different transverse and longitudinal relaxation rates in different parts of the tissue
4. Magnetisation transfer between water and a bound pool at membrane surfaces
5. Increased transverse and/or longitudinal relaxation when spins hit membrane surfaces
6. Surface tension causing spins to stick temporarily at membrane surfaces

**Table 1.** Properties of the tissue components. The properties marked by * can only be defined at the group-level. The others can be defined once for a whole group or for individual objects within the group.

| Parameter | Section | Description |
| --- | --- | --- |
| Type* | 2.1 | Wall, Cylinder, Sphere, or Triangle (part of Mesh) |
| Position/size | 2.1/2.3 | Determines location/size of the membranes. |
| Orientation* | 2.1/2.3 | Cylinder/wall orientation. Can be used to rotate mesh or a group of spheres. |
| Repeat scale* | 2.1/2.3 | On what length scale does the tissue geometry repeat itself |
| Magnetic susceptibility | 2.7 | Alters magnetic field strength experienced by spins in the neighbourhood around the object |
| **Collision parameters** |  |  |
| $\theta_{\text{relax}}$ | 2.9 | Sets relaxation rate |
| $\theta_{\text{perm}}$ | 2.10 | Sets membrane permeability |
| **Properties inside a tissue compartment (cylinder/sphere/mesh)** |  |  |
| $R_{1;\text{inside}}$ | 2.4 | Shift in the longitudinal magnetisation |
| $R_{2;\text{inside}}$ | 2.4 | Shift in the transverse magnetisation |
| **Bound pool parameters** |  |  |
| $\theta_{\text{bound}}$ | 2.11 | Surface density of bound pool |
| $\theta_{\text{transfer}}$ | 2.11 | Transfer time from bound to free pool |
| $R_{1;\text{bound}}$ | 2.4 | Shift in the longitudinal magnetisation |
| $R_{2;\text{bound}}$ | 2.4 | Shift in the transverse magnetisation |

The simulator supports any tissue geometry (represented by meshes) and any MRI sequence (represented as a series of radio-frequency (RF) pulses and gradient waveforms). It, therefore, is not limited to any specific tissue type and covers a wide range of MRI modalities.

The simulator is written in the Julia programming language, which allows both for easy interactivity and fast evaluation. In this manuscript, we describe the implementation of the simulator to construct the microstructural substrates and MRI sequences. Documentation and installation instructions can be found at https://michielcottaar.github.io/MRIBuilder.jl/stable/ for the sequence builder (MRIBuilder.jl) and at https://michielcottaar.github.io/MCMRSimulator.jl/stable/ for the simulator (MCMRSimulator.jl). All results were generated using V1.0 of the simulator.

## 2. Theory

In a real measurement, the MRI signal within a voxel is formed from the contribution of ∼10^20^ spins. Simulating all of these would be computationally prohibitive. In the simulator we use a Monte Carlo approach (Hall and Alexander, 2009), where we only simulate a representative subset of these to obtain an approximation of the true signal. This estimate of the true signal will contain some random noise, which will decrease as we increase the number of simulated spins.

Within the simulator each spin undergoes its own random walk restricted/hindered by the geometry of the simulated tissue. The magnetisation of each spin takes into account the MRI sequence and microstructural environment. Finally, the magnetisation is summed across all spins to generate the MRI signal. A more detailed overview of this algorithm is given in Figure 2 and the rest of this section.

**Figure 2:**
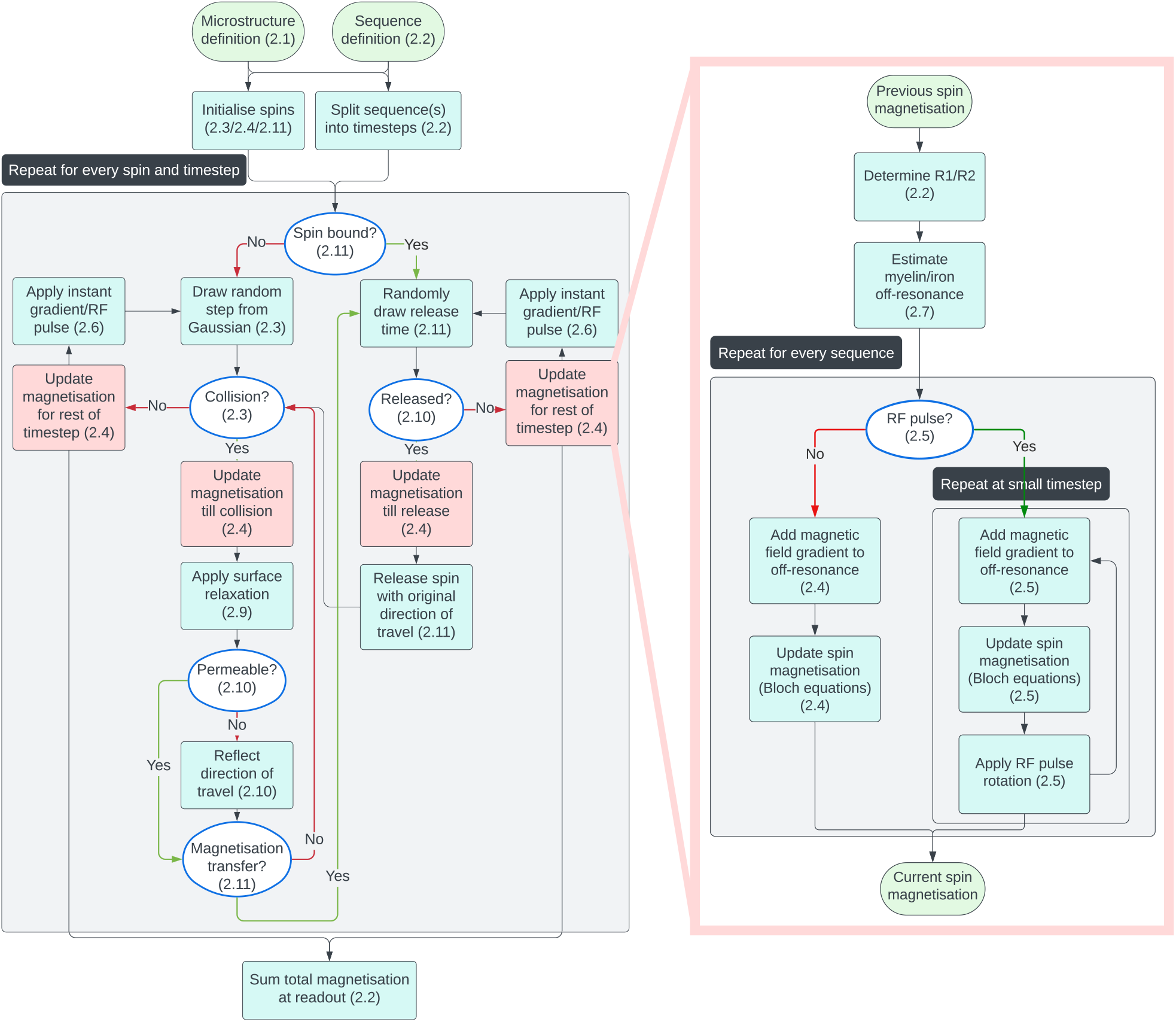
Flowchart of the Monte Carlo algorithm implemented in the MCMR simulator. User-defined sequence(s) and microstructure inputs (top) are initially processed. The remaining steps on the left are repeated for every spin and timestep. On the right the step to update the spin magnetisation (red boxes) is further expanded. For more details on each step check out the referenced section number.

Each spin represents a possible state history of the hydrogen nuclear magnetisation as it traverses the tissue. Spins are modelled as classical magnetic moments following the Bloch equations rather than quantum objects, which means that they actually represent a large group of hydrogen nuclear magnetisations with a similar trajectory (i.e., isochromats). For simplicity, we will still use the term “spin” throughout this manuscript.

The first two sections (2.1 and 2.2) deal with the main user inputs into the simulator, namely the geometry of the simulated tissue and the MRI sequence(s). The remaining sections contain details on how the MRI signal is computed.

### 2.1. Microstructure definition

One of the main inputs to the simulator is a computational representation of the tissue microstructure. This consists of any combination of walls, cylinders, spheres, or meshes.

Such representations of tissue microstructure can be obtained based on 3-dimensional electron microscopy images or based on generative computation models (Ginsburger *et al*., 2019; Palombo, Alexander and Zhang, 2019; Callaghan *et al*., 2020; Kleinnijenhuis *et al*., 2020; Villarreal-Haro *et al*., 2023).

Within the simulator, the microstructure model can affect the predicted MRI signal in number of ways:

1. It constrains the random walk of the spins due to collisions with the tissue membranes (section 2.3).
2. The longitudinal (T1) and transverse (T2) relaxation times can be different inside a tissue component versus outside (section 2.4).
3. Tissue with non-zero magnetic susceptibility alters the local magnetic field strength (section 2.7).
4. Spins colliding with the tissue might lose part of their magnetisation due to surface relaxation (section 2.9).
5. Spins might pass through a permeable membrane (section 2.10).
6. Spins can be bound to the tissue membranes after magnetisation transfer (section 2.11).

To control these effects, a wide variety of parameters can be set while constructing the geometry, which will be discussed in more detail in their respective sections (see Table 1). These parameters can be split into surface and volumetric parameters. Surface parameters (e.g., permeability, magnetic susceptibility, bound pool properties) can be set for each individual surface elements (e.g., mesh triangles). Volumetric parameters (i.e., the R1 and R2 inside a given tissue compartment) can be set separately for each individual cylinder/sphere or for each connected component of the mesh.

### 2.2. Sequence definition

The simulator can model the signal evolution for arbitrary sequences built out of any combination of RF pulses and gradient waveforms. RF pulses and gradient waveforms can be instantaneous or have a finite duration.

The sequences can be provided using the Pulseq file format (Layton *et al*., 2017), which means that the sequences can be defined in any software that supports this file format as output (e.g., pypulseq in python or pulseq in matlab). Sequences defined in the Pulseq file format can be run on MRI scanners across multiple vendors, which also means that an identical sequence can be used in both the simulator and for experimental data acquisition.

Alternatively, we provide a sequence builder tool (MRIBuilder.jl) that can be used to define sequences for the MCMR simulator. In brief, this tool allows one to define an MRI sequence as a series of RF pulses and gradient waveforms. Rather than setting the parameters that characterise these components directly, the user instead constrains any properties of the sequence (e.g., maximise the diffusion-weighted b-value for a fixed echo time of 80 ms). The parameters characterising the individual sequence components are then estimated using a non-linear optimiser (IPOPT, (Wächter and Biegler, 2006)). Sequences produced using this builder can also be written out in the Pulseq file format.

For computational efficiency, the simulator can predict the signal for multiple MRI sequences in a single run. When multiple sequences are passed on, the simulator will still only run a single Monte Carlo simulation during which the spin magnetisation is modelled independently for each sequence. This means that for each sequence the spins will have identical spatial trajectories, but a different magnetisation evolution.

The main output of the simulator is the total spin magnetisation summed over all spins during an instantaneous readout event. A continuous readout is approximated as a series of instantaneous readout events.

### 2.3. Simulating spin trajectory

The spin trajectory is given by a random walk constrained by collisions with any user-defined tissue representation. Given a tissue representation, for each spin the simulator will take the following steps during each timestep from *t* to *t* + τ (Figure 3A):

**Figure 3.**
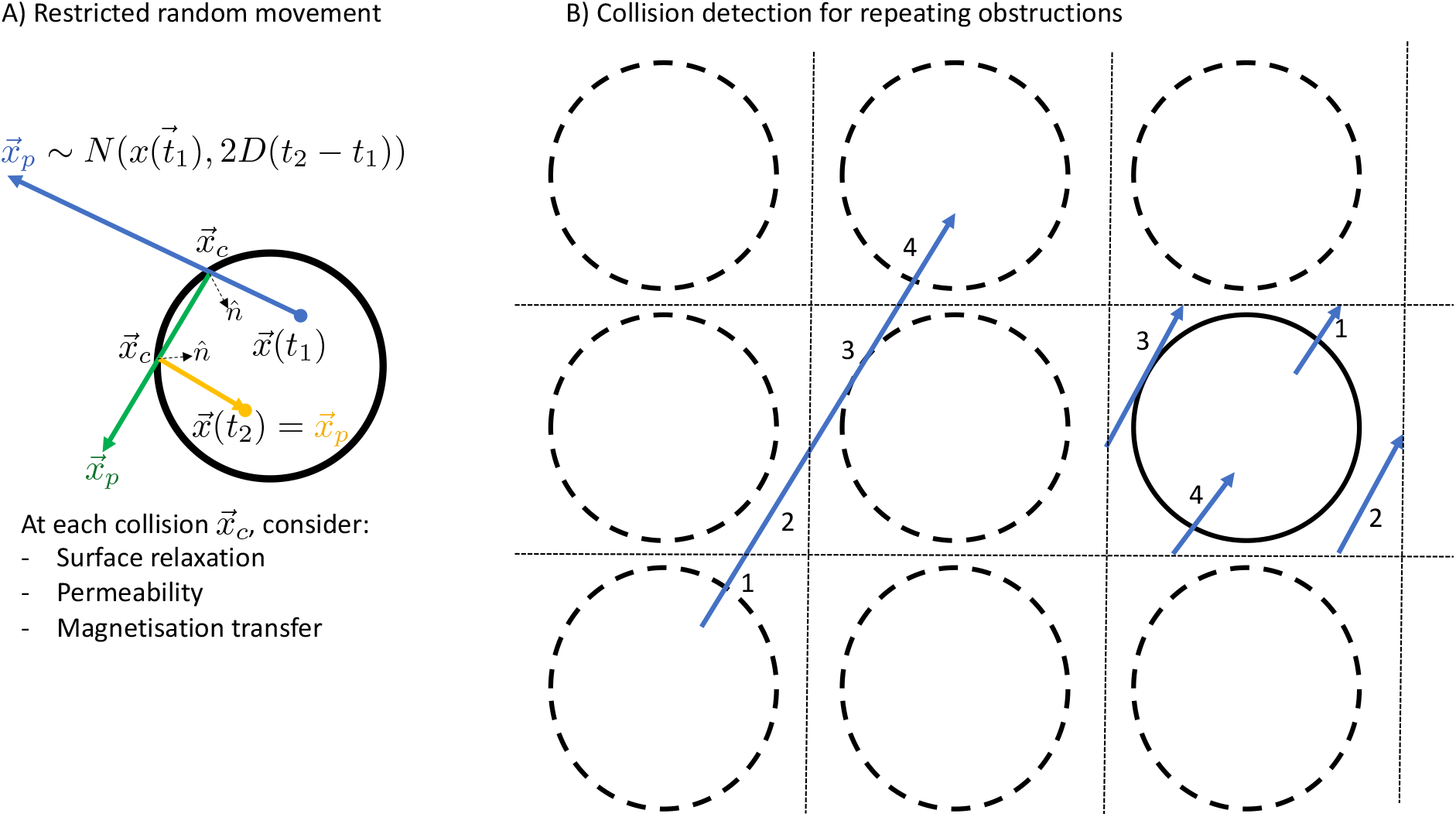
A) Illustration of the algorithm to model the trajectory of a spin between times t_1_ and t_2_ for a spin inside a cylinder. The initial displacement is drawn from a Gaussian distribution with variance 2D(t_2_ − t_1_) (blue). In this case, the spin undergoes two collisions in order to remain within the impermeably cylinders. At each collision the trajectory is reflected around the normal 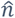 (green and orange). B) Illustration of a repeating geometry for a single cylinder (solid line), which is repeated on a grid (dashed lines). Each spin trajectory (e.g., left blue line) is projected onto the grid square (or cube) containing the actual cylinder. During this projection the trajectory will typically be split into multiple parts (labelled here as 1 to 4), which will be checked for collisions in sequence. Note that the diffusion step size illustrated in this plot is far larger than that used in the simulator.

1. Propose a new position ***x***_*p*_ from a 3D Gaussian distribution centred on the current position ***x***(*t*) with a variance 2*D*τ, where *D* is the diffusion coefficient:

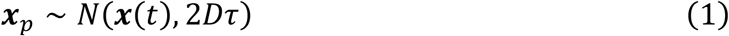
2. Identify the first (if any) collision ***x***_***c***_ between the line from ***x***(*t*) to ***x***_*p*_ and the user-provided geometry. If there is a collision, the default behaviour is to propose a new ***x***_*p*_′ by mirroring the trajectory around the normal 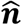 of the geometry (Figure 3A). Note that this behaviour may differ when modelling permeability or magnetisation transfer. We then repeat this step checking for any further collisions between ***x***_***c***_ and ***x***_*p*_′.
3. When there are no more collisions, we have our new position:

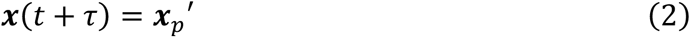

Any tissue representation will by necessity only have a limited spatial extent, which is typically smaller than the MRI field of view (or even a single MRI voxel). To simulate the signals from larger volumes, the microstructure within the simulator can repeat itself on a user-defined length scale. To detect collisions in such a repeating geometry, we take the modulo of the spin trajectory with respect to the repeating length scale (Figure 3B).

For computational efficiency reasons, we pre-compute the intersections of the obstructions with a grid. When a spin passes through a grid element, we then only have to consider possible intersections with those obstructions rather than looping over all obstructions.

Within the simulator the timestep is determined in an automatic manner as described in supplementary material A. This timestep is optimised to obtain accurate results in a computationally efficient manner.

### 2.4. Updating spin magnetisation

While spins traverse the geometry as described above, their magnetisation is updated according to the Bloch equations. We keep track of both the transverse and longitudinal magnetisation. At any time in the sequence, the signal can be read out by summing the magnetisation vectors of all of the spins. The MRI signal is then obtained by taking the transverse component of the total magnetisation vector.

Whenever a collision is detected in the spin trajectory, the spin magnetisation is updated up until the time of the collision before processing the collision itself. This ensures that the spin magnetisation used during the collision is accurate, which is important for modelling magnetisation transfer and surface relaxation as discussed in later sections (Figure 2).

In the absence of an RF pulse, the Bloch equations can be analytically solved:

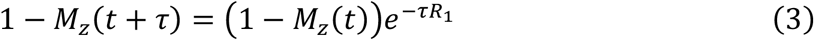

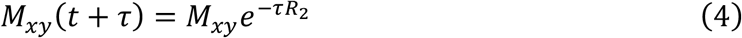

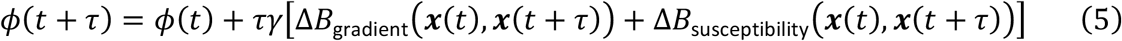

where *M*_*Z*_ and *M*_*xy*_ are the longitudinal and transverse magnetisations, ϕ is the phase of the spin magnetisation in the transverse plane, and τ is the timestep. *R*_1_ and *R*_2_ are the user-provided longitudinal and transverse relaxation rates, which might be different for spins in different local environments (Table 1). Note that the equilibrium magnetisation (*M*_*z*,0_) is assumed to be 1 (in arbitrary units) for every spin.

The Bloch equations are solved in the frame rotating with the Larmor frequency (γ*B*_0_). In this rotating frame the spin phase is only affected by deviations from the main magnetic field (Δ*B* in eq. 5). This off-resonance field has two contributors, namely the MRI magnetic field gradients (Δ*B*_gradient_) and the fields generated due to the tissue magnetic susceptibility (Δ*B*_susceptibility_). The latter is described in section 2.7.

The average MRI magnetic field gradient experienced by a spin in a single timestep can be calculated by solving the following integral

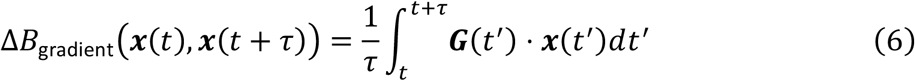

The timesteps are chosen in such a way that the MR scanner gradient is well approximated as changing linearly over time within each timestep (Supplementary Material A). For a spin travelling in a straight line from ***x***(*t*) to ***x***(*t* + τ), this leads to:

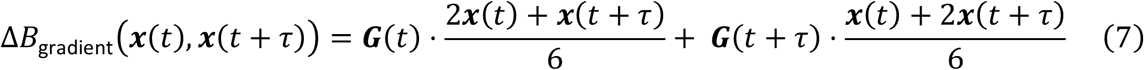

### 2.5. Applying RF pulses

During an RF pulse, the simulator solves the Bloch equations by splitting each timestep further into very short chunks of duration τ_RF_. During each such chunk, the RF pulse is approximated as a hard RF pulse, which causes the spin magnetisation vector to rotate around the following vector:

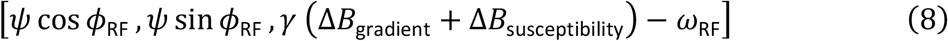

The angle of this rotation is given by the norm of this vector. *ψ* is the flip angle of the RF pulse, which is given by the integral of the RF pulse amplitude over this chunk of time. ϕ_RF_ and *ω*_RF_ are, respectively, the phase and off-resonance frequency of the RF pulse. After each rotation step, relaxation is applied as described in equations 3-5. The rotation step above is accurate in the coordinate system rotating with the RF pulse (in which the orientation of the RF pulse can be assumed to be constant). To get the resulting spin magnetisation back into the coordinate system rotating with the Larmor frequency, we add *ω*_RF_τ_RF_ to the spin’s phase.

The rotation and relaxation steps above get repeated many times during a single timestep. Each repetition has a duration of τ_RF_, where τ_RF_ is guaranteed to be short enough that the RF pulse amplitude and frequency can be assumed to be roughly constant and short compared with the spin’s *T*_1_ and *T*_2_. During each of these repetitions we iteratively apply the RF pulse and allow for relaxation. The gradient contribution to the off-resonance field (Δ*B*_gradient_) is re-evaluated in each iteration, but for efficiency reasons the susceptibility component (Δ*B*_susceptibility_) is only evaluated once (far right in Figure 2). The overall timestep is set to be short enough for such a single evaluation of the susceptibility component to give accurate results (Supplementary Materials A).

### 2.6. Applying instantaneous RF pulses and gradients

While not realistic, instantaneous approximations of RF pulses and MR gradients are often considered when discussing sequences or analytically deriving sequence properties. To validate such analytic derivations as well as enable comparisons between idealised and more realistic versions of a sequence, we allow instantaneous RF pulses and gradients to be used in the simulator.

Given our definition of the timesteps (Supplementary Materials A), these instantaneous components are always guaranteed to be at the edges between timesteps. At these times the spins have a well-defined location and magnetisation (Figure 2), which allows for straightforward implementations of these components.

Instantaneous RF pulses are defined by their phase (ϕ_RF_) and flip angle. Because they are instantaneous, they have an infinite bandwidth, which means that it is not meaningful to talk about their off-resonance frequency. They are implemented through rotation of the spin magnetisation by the flip angle around the following vector:

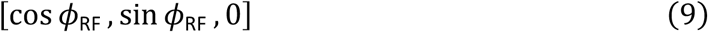

Instantaneous MR gradients are defined by the area under the curve or **q**-vector. The spin phase is updated based on the spin’s current location ***x***:

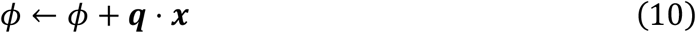

### 2.7. Off-resonance fields

Many constituents of biological tissue do not have the same magnetic susceptibility as water. This means that when placed in the strong magnetic field of an MRI scanner, the tissue will change the local magnetic field strength, which causes dephasing of the MRI signal. The simulator models the contributions to these local magnetic field perturbations from cylindrical configurations or arbitrary meshes. Both cylinders and meshes can have isotropic and/or anisotropic magnetic susceptibility.

Figure 4 illustrates the off-resonance fields produced by a myelinated annulus (i.e., nested cylinders), a cylinder, and a single triangle within a larger mesh. For the myelinated annulus we use the hollow fibre model as given by Wharton and Bowtell (2012) (Figure 4A). We also allow single cylinders to be “myelinated”, even though there is no actual room for the myelin. In this case we still use the equations from Wharton and Bowtell (2012) for the intra- and extra-axonal space for a user-provided *g*-ratio (i.e., inner over outer radius of the myelin) without actually having a myelin sheath taking up any physical space (Figure 4B).

**Figure 4.**
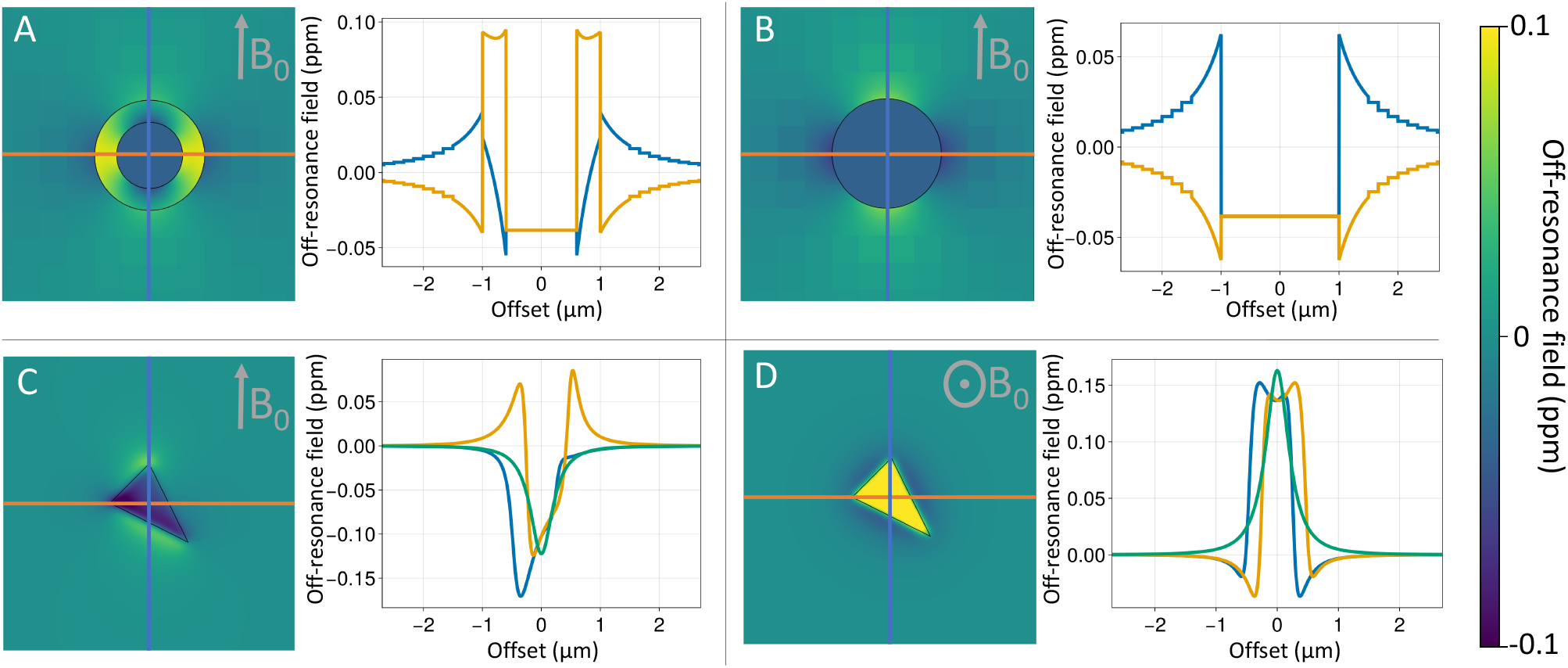
A) Off-resonance fields produced by the hollow-fibre model for myelinated axons, (B) an approximation of this hollow-fibre model that ignores the physical space actually taken up by the myelin and (C, D) a single triangle (which could be part of any larger mesh). In each panel the off-resonance field is shown on the left, with the line plots showing the off-resonance field along the 3 primary axes (blue and orange in-plane and green out of plane).

This is a non-physical model that allows one to approximate the effect of the magnetic susceptibility of myelin on the intra- or extra-axonal space without considering its geometric effect (or the contribution of myelin water).

The hollow fibre approximation assumes infinitely long, perfectly straight cylinders. In reality, axons are bendy as they weave themselves through the brain’s white matter. Moreover, myelin sheaths are separated by short gaps at the nodes of Ranvier and many axons alternate between myelinated and unmyelinated segments along their length. To support this rich variation in geometry we the simulator is able to calculate the off-resonance field from arbitrary meshes. We do this by summing over the contribution from each mesh triangle, with each contribution being calculated as described in Rubeck *et al*., (2013) (Figure 4C,D).

Computing the off-resonance field experienced by a particle travelling between positions ***x***_1_ and ***x***_2_ is computationally challenging as it involves integrating the contribution from each magnetic susceptibility source over all possible trajectories between ***x***_1_ and ***x***_2_. We approximate these integrals by just evaluating the field at a single random point along the line connecting ***x***_1_ and ***x***_2_ and make the timestep short enough that this approximation holds (Supplementary Materials A).

A key challenge is that the effect of magnetic susceptibility is “non-local” (see Figure 4) and explicitly summing over the contribution from each individual cylinder or mesh triangle for every spin location would be computationally very expensive. Instead, we overlay a grid on the geometry and compute the off-resonance field in the following way:

1. For geometry elements close to a given spin we directly compute their contribution to the off-resonance field as described above.
2. For more distant geometric elements we pre-compute their contribution on the grid and add this total to the spin’s off-resonance field. This assumes that the field contribution is constant over a single grid element, which will only be true if the grid element size is small compared with the distance from the geometric element. We apply this approximation only for contributors more than 3 grid element sizes away.
3. For a mesh, we define an intermediate group, where the contribution of triangles that are more than 4 times the size of the triangle away from the spin are approximated as dipoles rather than using the more computationally expensive equations for a triangle (Rubeck *et al*., 2013). This approximation is applied for cylinders and annuli as well, but this has no effect on the result because the dipole approximation is exact for these configurations.

### 2.8. Timestep dependent collision rate

A complication in implementing the simulator is that the constrained random walk resulting from the algorithm described in 2.3 is only a rough approximation of the far more convoluted path the spins might take in reality. As long as the timestep is short enough, ideally this approximation would not affect any properties that the MRI signal is actually sensitive to. However, this is not the case for the number of collisions. While this timestep dependence of the collision rate is unavoidable, the simulator is designed in such a way that this timestep dependence does not actually affect rate of the phenomena that happen at these collisions, such as surface relaxation, permeability, and magnetisation transfer. We discuss how this is achieved in the upcoming sections discussing these phenomena (2.9-2.11). These corrections are as far as we know derived here for the first time and are crucial to accurately model surface-based MR effects in a Monte Carlo simulator.

A single collision in the simulator represents many collisions along the convoluted path that the spin would have taken in reality. The reason for this can be appreciated by considering that within a single timestep the number of collisions is proportional to the mean displacement ⟨*x*⟩ and, hence, the square root of the timestep τ:

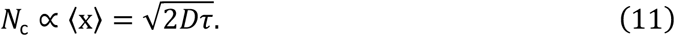

While the number of collisions per timestep increases for larger timesteps, the rate of collisions actually decreases as the timestep increases:

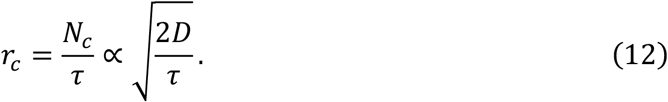

This means that if we run the same simulation twice, once with a timestep of 0.1 ms and once with a timestep of 1 ms, the first simulation will have 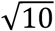 times more collisions than the second.

To relate these collision-specific behaviour with more biophysical parameters, we derive in S2 the full equation for the collision rate of a spin within a volume *V* enclosed by a surface *S*:

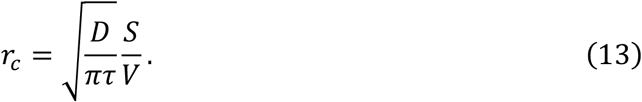

This equation will be used in later sections to compute parameters like the exchange rate or effective relaxation rate from the simulator parameters.

### 2.9. Surface relaxation

Spins close to a membrane often relax more quickly than spins further away from membranes due to a combination of multiple effects including reduced spin tumbling and magnetisation transfer (Schyboll *et al*., 2019). While this can be partially modelled using a bound pool (2.11), we also allow direct simulation of surface relaxation. This is implemented by reducing the spin magnetisation at each collision:

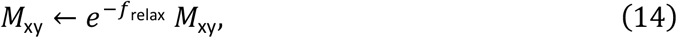

where *M*_xy_ is the transverse magnetisation, and *f*_rela*x*_ is a positive number representing the rate with which the magnetisation relaxes at each collision.

To ensure that the resulting relaxation rate does not depend on the timestep of the simulator, we need to compensate for the fact that the collision rate does depend on the timestep τ (2.8). To derive this, we will consider the attenuation rate due to the surface relaxation after some time *t*. During this time, we expect *r*_*c*_(τ)*t* collisions resulting in a signal attenuation of:

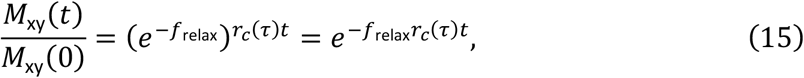

Thus, the only way to ensure that the resulting attenuation is independent of the timestep of the simulator is to make *f*_relax_ depend on the timestep. We do this by defining another timestep-independent parameter *θ*_relax_ which is related to *f*_relax_ as follows:

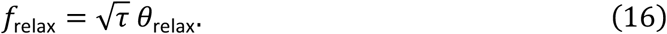

When defining a rate of surface relaxation, the user actually sets *θ*_rela*x*_ rather than *f*_relax_.

Within a closed surface the resulting relaxation rate can be computed analytically using the equation for the collision rate (eq. 13):

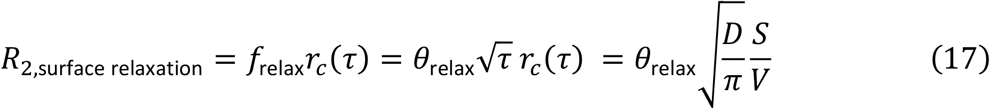

### 2.10. Permeability

The default behaviour for spins hitting the membrane is to reflect their direction of travel around the membrane surface normal (Figure 3A), however in reality water might also pass through a membrane (either directly or through cross-membrane channels or pumps). To model this, we allow users to set the permeability of individual objects within the tissue microstructure. If the spin collides with a permeable object, it might pass through the object with a given probability or be reflected.

To ensure that the exchange rate does not depend on the simulation timestep (2.8), we define the user-set parameter *θ*_perm_ in such a way that the exchange rate of spins from compartment A to B is:

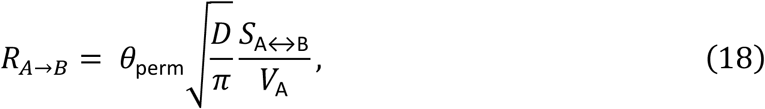

where *S*_A↔B_ is the surface area of the membrane between compartments A and B and *V*_A_ is the volume of compartment A.

This parameter *θ*_perm_ is related with the probability of spins passing through the membrane by:

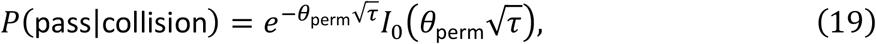

where *I*_0_ is the zero-order modified Bessel function of the first kind. This equation can be derived by considering that the fraction of spins retained in the original compartment has to be constant as the timestep increases (Sukstanskii, Yablonskiy and Ackerman, 2004). If spins were unable to return to the original compartment, we would only have the exponential term. The Bessel function is added to correct for the decreased probability of spins returning to the original compartment as the timestep increases (Figure S2).

Even with this correction the simulator will still inaccurately model exchange if the timestep is too long, because within a single timestep a particular spin can only pass through the membrane zero or one times. However, if the timestep is too long a typical spin might in reality have passed through the membrane many times. To avoid this situation from arising, we keep the timestep short enough, so that *P*(pass|collision) is much lower than one. This is achieved when:

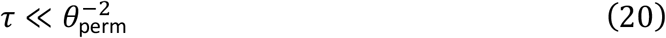

A special case is a perfectly permeable surface represented by *θ*_perm_ = ∞. Such a surface has no effect on the spin trajectory and is implemented by setting *P*(pass|collision) to 1 with no constraint on the timestep. Note that we still detect and process collisions with such permeable surfaces in order to model other effects such as surface relaxation, magnetisation transfer, and/or the transition of the spins between different compartments (with different relaxation properties).

#### 2.11. Magnetisation transfer

Magnetisation transfer is modelled by the inclusion of a bound pool and the possibility of spins to transition between the bound and free pool (i.e., chemical exchange or dipole-dipole interactions). In the simulator the bound pool can only be localised to the membranes (i.e., the user-provided tissue boundaries). This creates a fundamental difference from many descriptions of magnetisation transfer where a spin from the free pool can join the bound pool at any time and location. In our simulator a free spin can only join the bound pool at the time of a collision.

Magnetisation transfer is characterised by two parameters, which can be set by the user independently for each object (Table 1). The first of these is the density of the bound pool relative the free pool (*θ*_bound_). This is defined as the ratio of the surface density of the bound pool (Σ_bound_) over the volume density of the free pool (*ρ*_free_):

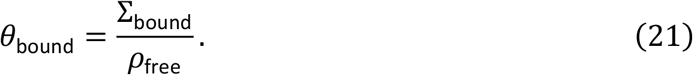

This parameter has units of μm. It can be thought of as the distance that you have to travel away from the surface to see the same number of free spins as those bound to the surface. If the surface-to-volume 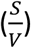 ratio of the tissue is known, it can be used to compute the fraction of spins in the bound pool:

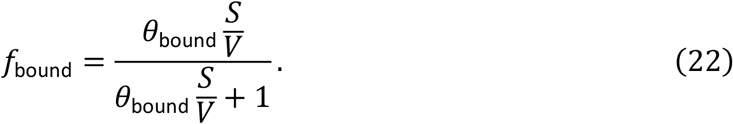

During spin initialisation, spins are assigned to the bound pool with random positions on the tissue membranes until the correct surface density is achieved.

The second parameter sets the timescale at which individual spins transfer from the bound to the free pool (*θ*_transfer_). The time until release for a bound spin is drawn from an exponential distribution:

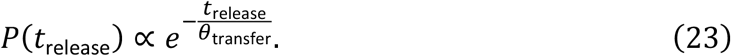

This distribution ensures that the probability of a spin being released at any point does not change over time. *θ*_transfer_ is the average dwell time of a bound spin before it is released into the free pool.

We can estimate the probability that a spin should transfer to the bound pool at a collision based on the two parameters defined above. This derivation uses the fact that the rate of spins going from the free to the bound pool should match the rate of spins going from the bound to the free pool:

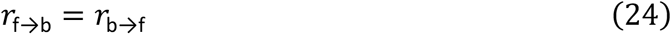

The rate from the free to the bound pool is given by:

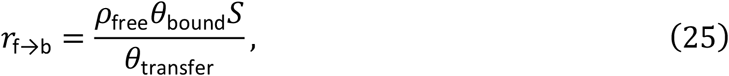

where the numerator is the total number of bound spins over a surface *S* and the denominator is the average time each bound spin spent being bound.

The rate from the bound to the free pool is given by:

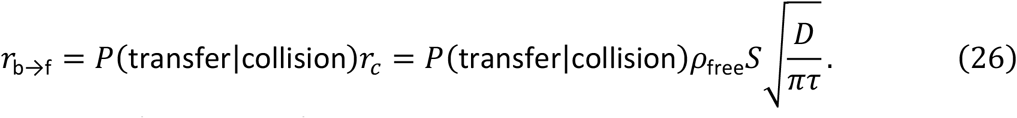

Solving these equations for the transfer probability at a collision gives:

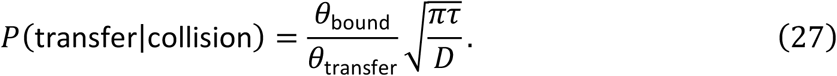

While the equation above is accurate for small timesteps, it breaks down for larger timesteps, when this equation for *P*(transfer|collision) starts approaching or even exceeding 1. We find the transfer rate to be more robust at long timesteps when adopting (Figure S2):

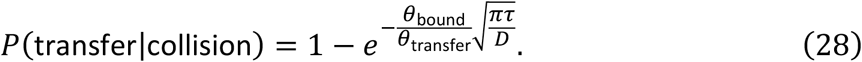

As we have argued for the permeability, the probability of magnetisation transfer depends on the simulation timestep. A timestep that is too large will have a collision rate lower than the transfer rate (*r*_b→f_) and, hence, give an unphysical result of P(transfer|*c*ollision) > 1. In order to ensure that the simulation contains both collisions with and without magnetisation transfer, we add an upper limit to the timestep to be ensure that P(transfer|*c*ollision) ≪ 1:

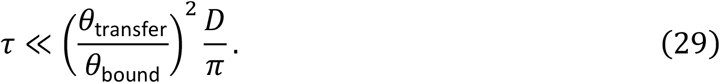

## 3. Results

We illustrate the use of the simulator for four commonly used MRI sequences, namely diffusion-weighted MRI, magnetisation transfer imaging, spin echo, and gradient echo. We will model the effect of all the different features of the simulator discussed above. The results presented here are selected to illustrate the flexibility of the simulator and to validate the simulator in comparison with analytical models. The novelty here lies in the fact that the same simulation environment is capable of investigating a wide range of MR phenomena and contrast mechanisms. Sequences were plotted using the plotting support included in the simulator itself (based on Makie.jl; Danisch and Krumbiegel, 2021). The Jupyter notebooks used to create these plots are available at https://git.fmrib.ox.ac.uk/ndcn0236/mcmr_paper_figures.

### 3.1. Diffusion

Figure 5 shows the signal dependence of diffusion MRI on diffusion time for an idealised diffusion-weighted spin echo sequence with instantaneous RF pulses, gradients, and readout at a fixed b-value of 0.1 ms/μm^2^. The microstructure model is a set of randomly positioned infinite cylinders (diameter of 10 μm and packing density of 0.6) with the same orientation perpendicular to the diffusion gradient. At very short diffusion times, very few spins interact with the membranes leading to an attenuation of *e*^−*bD*^ ≈ 0.74. As the diffusion time increases we get a dependence of the signal attenuation on the S/V ratio as described by Mitra *et al*. (1992) for both intra and extra axonal spaces. At even longer diffusion times, the spins inside the axons effectively become trapped leading to no attenuation. At this low diffusion weighting (b=0.1 ms/μm^2^) this transition from free to restricted diffusion is very well described for the intra-axonal signal by the Gaussian phase approximation (van Gelderen *et al*., 1994) or the short gradient-duration approximation from Callaghan (1995) (latter not shown).

**Figure 5.**
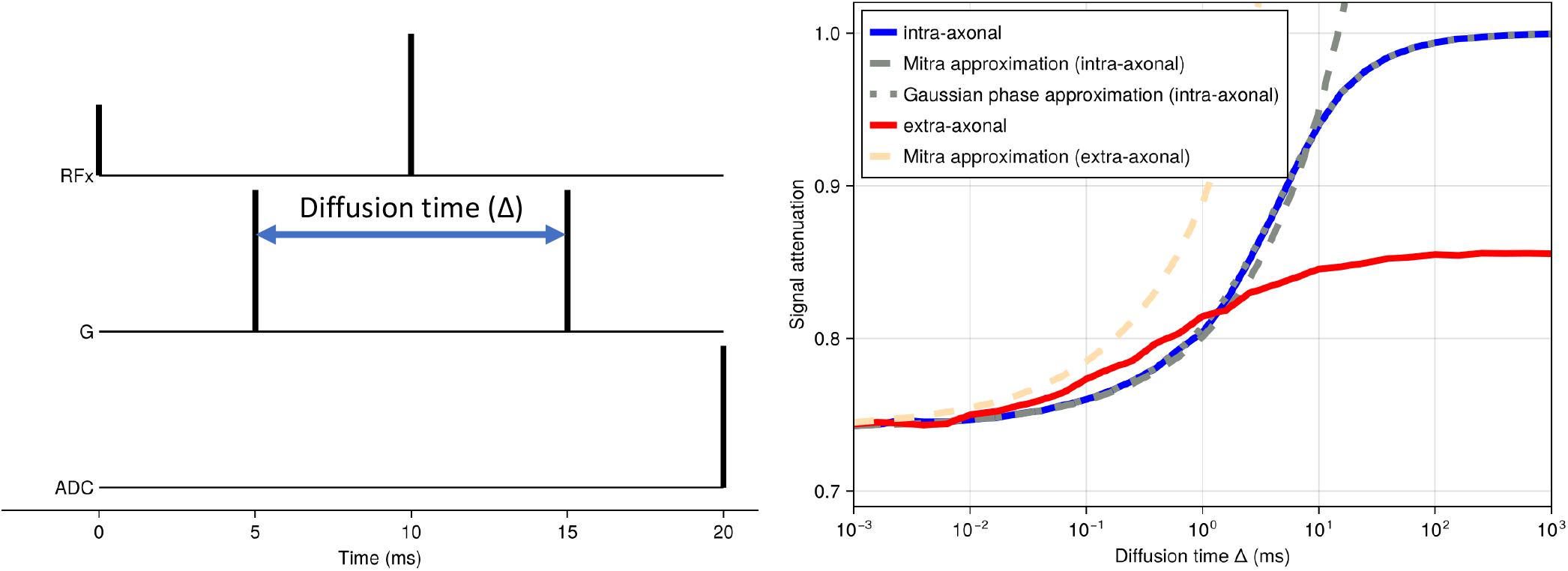
MCMR Simulator results for diffusion MRI sequences for a simplified model of the brain’s white matter, namely a set of randomly distributed parallel cylinders (i.e., “axons”) with a diameter of 10 μm and a packing density of 0.6. The intrinsic diffusivity is set to 3 μm^2^/ms. We investigate the diffusion MRI signal dependence on diffusion time in an idealised sequence with instantaneous RF pulses, gradients, and readout. Across all diffusion times the diffusion weighting is kept low at a b-value of 0.1 ms/μm^2^ with gradient orientations perpendicular to the cylinders. The signal is split into the intra-axonal (blue) and extra-axonal (red) components with analytical approximations overlaid as dashed lines for the (Mitra et al., 1992) approximation and as dotted line for the Gaussian phase approximation inside the cylinders (van Gelderen et al., 1994).

In addition to such idealised sequences, the simulator can also be used to model more realistic sequences. For example, Figure 6 shows the shortest sequence that can be obtained with a b-value of 20 ms/μm^2^, an isotropic voxel size of 2 mm, and an echo planar imaging (EPI) readout resolution of 60 by 40 (no acceleration) on a scanner with a maximum gradient 300 mT/m and a slew rate of 200 mT/m/ms (i.e., equivalent of Siemens Connectom scanner; Setsompop *et al*., 2013). The diffusion-weighting gradients are scaled to vary the effective b-value. The microstructural model is the same as for Figure 5. Under these conditions, the intra-axonal signal attenuation deviates significantly from the Callaghan approximation (Callaghan, 1995), which assumes short gradient durations. The Gaussian phase approximation (van Gelderen *et al*., 1994) does still provide a reasonable match.

**Figure 6.**
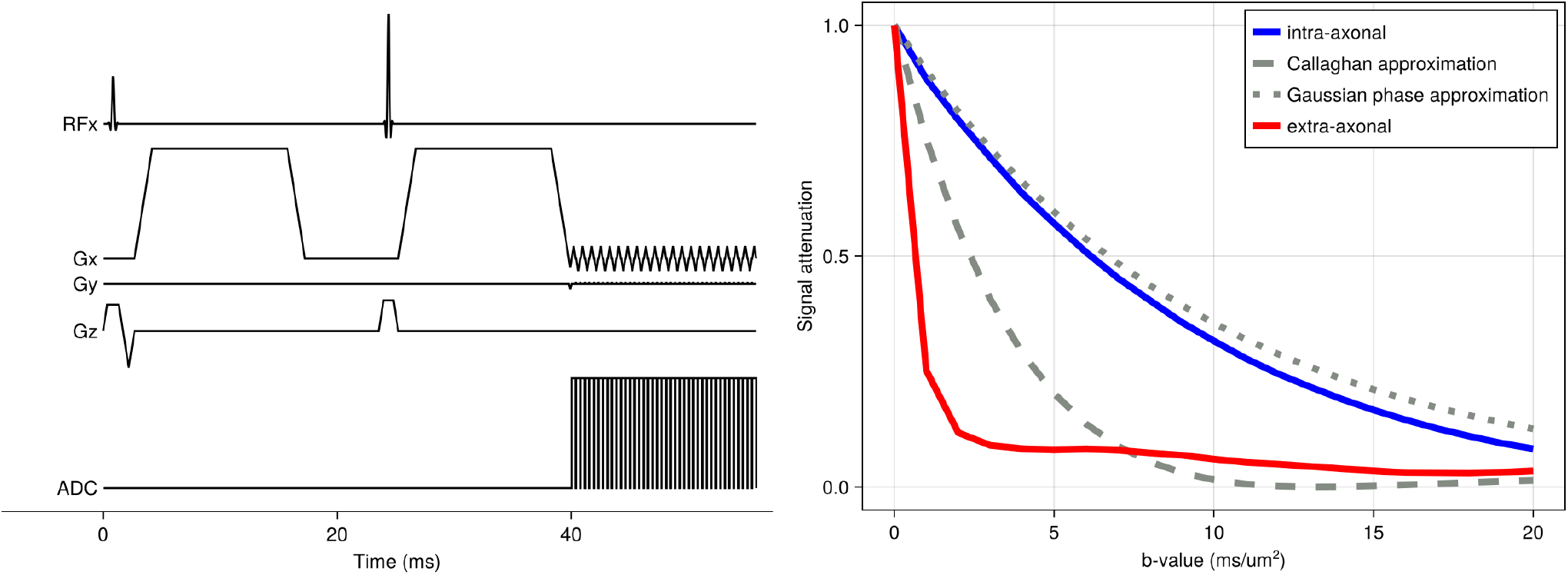
MCMR Simulator results for diffusion MRI sequences for a simplified model of the brain’s white matter, namely a set of randomly distributed parallel cylinders (i.e., “axons”) with an intrinsic diffusivity of 3 μm^2^/ms. We investigate the diffusion MRI signal dependence on the strength of the diffusion weighting (b-value) in a realistic sequence with slice-selective RF pulses, gradients, and EPI readout (δ=13 ms; Δ=23 ms). The signal is split into the intra-axonal (blue) and extra-axonal (red) components. For the intra-axonal component analytical approximations are overlaid based on the Gaussian phase approximation (van Gelderen et al., 1994) and the short-gradient duration approximation (Callaghan, 1995).

### 3.2. Magnetisation transfer

While previous Monte Carlo simulators of the tissue microstructure have focussed on diffusion-weighted MRI, MCMR simulator can be used to investigate alternate contrast mechanisms and associated sequences. As an example, we here illustrate the saturation pulse used in a magnetisation transfer sequence.

The saturation pulse consists of regular off-resonance pulses (top in Figure 7), which saturates the magnetisation in the bound pool (dashed lines in Figure 7). The reason the spins in the bound pool are affected like this is that they have been assigned a short T_2_ of 0.1 ms (much shorter than the 80 ms for the free pool). There are two effects of this short T_2_, both of which are captured by the simulator. First, the short T_2_ increases the bandwidth of the bound pool, which means that it is more affected by the off-resonance pulse than the free pool. This linewidth broadening is captured in the simulation by explicitly solving the Bloch equations during the RF pulse without having to be explicitly implemented. Second, the short T_2_ means that any transverse magnetisation excited by the pulse is immediately lost, which leads to the net saturation of the magnetisation of the bound pool.

**Figure 7.**
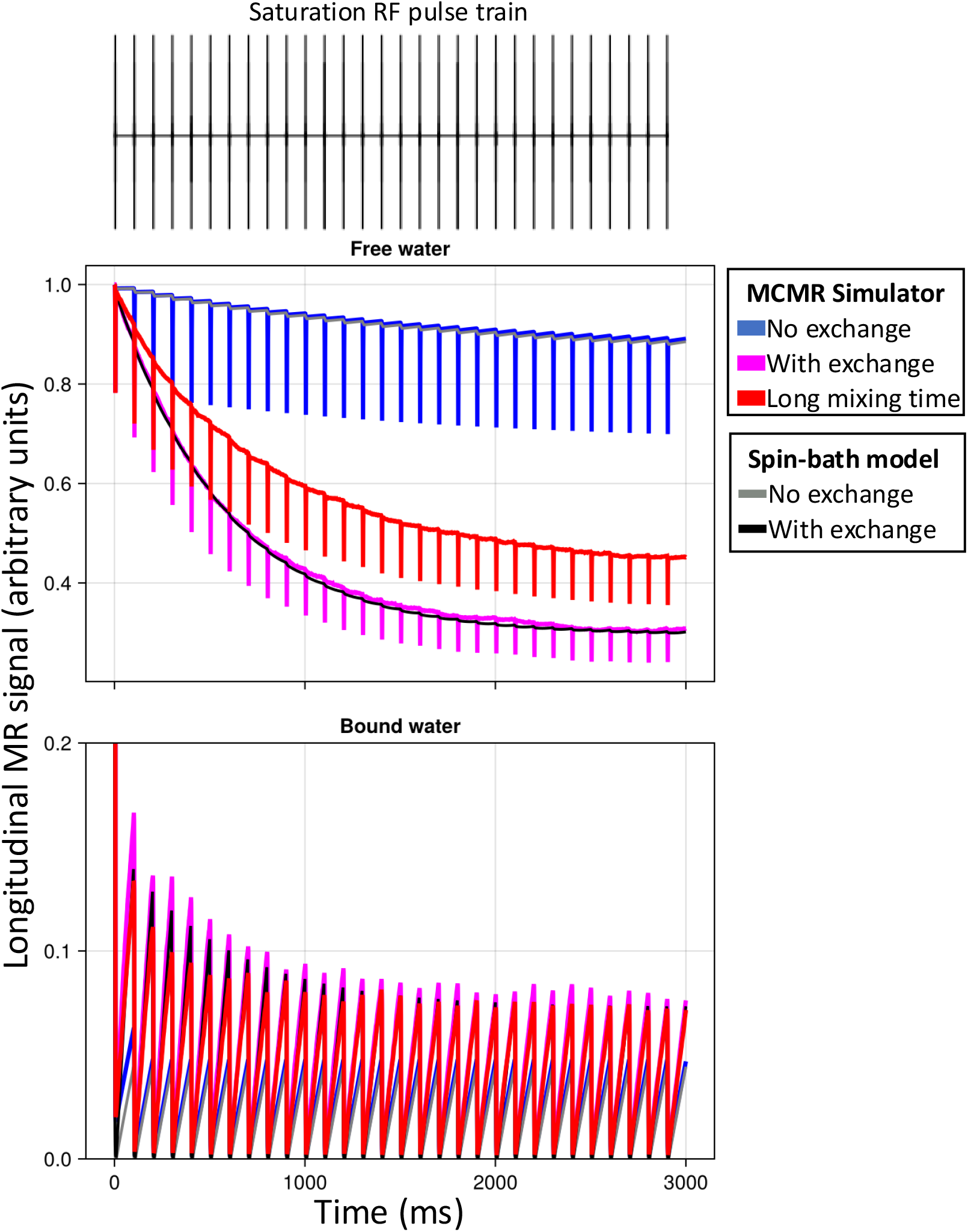
The longitudinal magnetisation evolution for the free (top) and bound (bottom) pool during a saturation pulse under three conditions. The saturation pulse consists of 30 sinc RF pulses with a duration of 10 ms, a flip angle of 700°, and an off-resonance frequence of 1.5 kHz (as recommended by De Boer (1995)). Half of the spins are in the bound pool; both pools have a T_1_ of 2 seconds. The free pool has a T_2_ of 80 ms and the bound pool of 0.1 ms. The microstructure consists of repeating, infinite walls every 1 μm with the bound pool being bound to those walls. This configuration ensures that there is, in effect, only a single free water pool. Blue and magenta show the evolution of the longitudinal magnetisation without and with magnetisation transfer (with a transfer time of 0.8 seconds). In both these scenarios, the free water diffusivity is 3 μm^2^/ms, which ensures that the free compartment is effectively mixed (mixing time on the order of 1 ms), which causes these results to closely match that of the spin-bath model from (Henkelman et al., 1993) (in orange and cyan). In the red scenario, the diffusivity is greatly decreased to 10^−5^ μm^2^/ms, so that the free-water mixing time increases to about 1 s (note that in the simulations this reduction in diffusivity does not affect the magnetisation transfer rate between the bound and free pool). This reduced mixing cannot be captured by the spin-bath model.

If we do not simulate any magnetisation transfer between the bound and free pool (blue in Figure 7), then this saturation of the bound pool will not affect the free pool and we can only detect the small, direct effect that the off-resonance pulses have on the free pool magnetisation. So, the magnetisation transfer ratio (i.e., ratio of the free water magnetisation with and without this saturation pulse) is small in this case. As we add magnetisation transfer (magenta) we can see a larger reduction in the longitudinal magnetisation of the free pool as detected in magnetisation transfer experiments. Traditionally, the size of this reduction is modelled using the spin-bath model (Henkelman *et al*., 1993), which assumes a dependence on the size of the bound pool and the transfer rate between the pools. Our simulations however also show a dependence on the mixing of the free pool, as is illustrated by lowering the diffusivity (i.e. increasing mixing time while keeping the magnetisation transfer rate constant) (red).

### 3.3. Relaxation

Finally, we investigate how axon properties, such as myelin thickness or radius, might affect the effective T_2_ from a spin echo sequence or T_2_* from a gradient echo sequence (Figure 8). The goal is to illustrate how the complex behaviour captured by the simulator causes these effective T_2_ and T_2_* to deviate from the true value (set to 2 seconds as observed in the cerebrospinal fluid in the ventricles).

**Figure 8.**
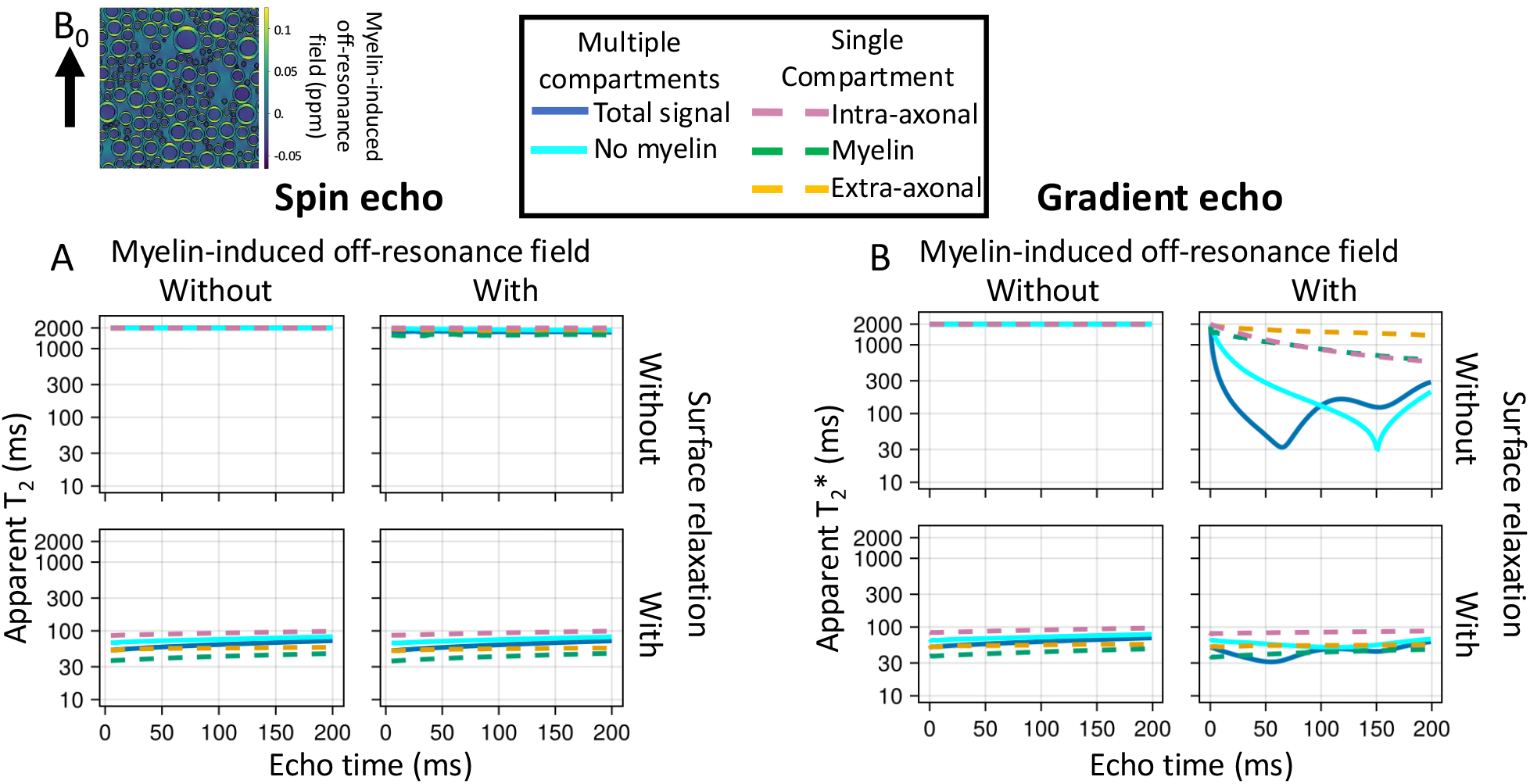
Simulations of a spin echo (left) or gradient echo (right) sequences for myelinated axons. The sequences are very simple and consist of an instantaneous excitation pulse, an instantaneous refocus pulse at half the echo time (only for spin echo), and an instant readout at the echo time. The myelinated axons are represented by 2 impermeable cylinders at the inner and outer radius of the myelin sheath (upper left), which effectively splits the water into three compartments (34% intra-axonal in magenta dashed line; 36% myelin in green; and 30% extra-axonal in yellow). The signal summed over these compartments are also plotted as blue, solid lines. The axons are all orthogonal to the main magnetic field (B_0_). The ratio of the inner to outer radius (i.e., g-ratio) is kept constant at 0.7. A, B) The trend of T2 (left) and T2* (right) as a function of echo time without or with a myelin-induced off-resonance field (isotropic susceptibility χ_I_ and anisotropic susceptibility X_A_ both set to -0.1 ppm, Wharton and Bowtell (2012)) and without or with surface relaxation (i.e., signal loss at every collision with the membranes).

In Figure 8 we investigate two potential contributors, namely the myelin-induced off-resonance field and surface relaxation. The off-resonance field has little effect on the T_2_ measured from a spin echo sequence as its effect is nearly perfectly cancelled out by the refocusing pulse. However, the magnetisation lost through surface relaxation is non-recoverable, leading to lower T_2_. The amount of surface relaxation was chosen to match the actual T_2_ measured in brain tissue of about 70 ms, which effectively makes the assumption that the actual T_2_ is dominated by the surface relaxation.

Because the different compartments have different surface-to-volume ratios and, hence, different collision rates, the single surface relaxation value does result in different T_2_’s for each compartments (∼80 ms for intra-axonal, 60 ms for extra-axonal, and 45 ms for myelin). These intra- and extra-axonal values are consistent with those observed in brain tissue at 3T, despite being controlled by only a single global variable (Veraart, Novikov and Fieremans, 2018; McKinnon and Jensen, 2019). While the myelin T_2_ is far too long (Lee *et al*., 2021), this might reflect that the simplified geometry of an inner and outer cylinder adopted here is a poor representation of the myelin water’s environment (Canales-Rodríguez *et al*., 2024).

In the gradient echo simulations, adding a myelin-induced off-resonance field does have a large effect on the T_2_* (solid lines on right in Figure 8B), although not within individual compartments (dashed lines). This is because each compartment is well mixed due to diffusion such that each spin experiences a magnetic field that is close to the average of that compartment. However, this average magnetic field differs between compartments, causing the compartments to accumulate phase differences due to precession. As a result, the different compartments come in and out of phase with each other over time and the T_2_* is substantially lower for the signal summed over compartments (solid lines on right in Figure 8B).

Figure 9 illustrates how microstructural properties affect the T_2_ and T_2_*. The effect of surface relaxation on the T_2_ is mainly mediated through the surface-to-volume ratio. Hence, the T_2_ in these simulations increases if we increase the axon diameter (in all compartments, Figure 9A) or reduce the axon density (in the extra-axonal compartment, Figure 9B). Such a T_2_ dependence on axon diameter has been reported previously (Barakovic *et al*., 2023).

**Figure 9.**
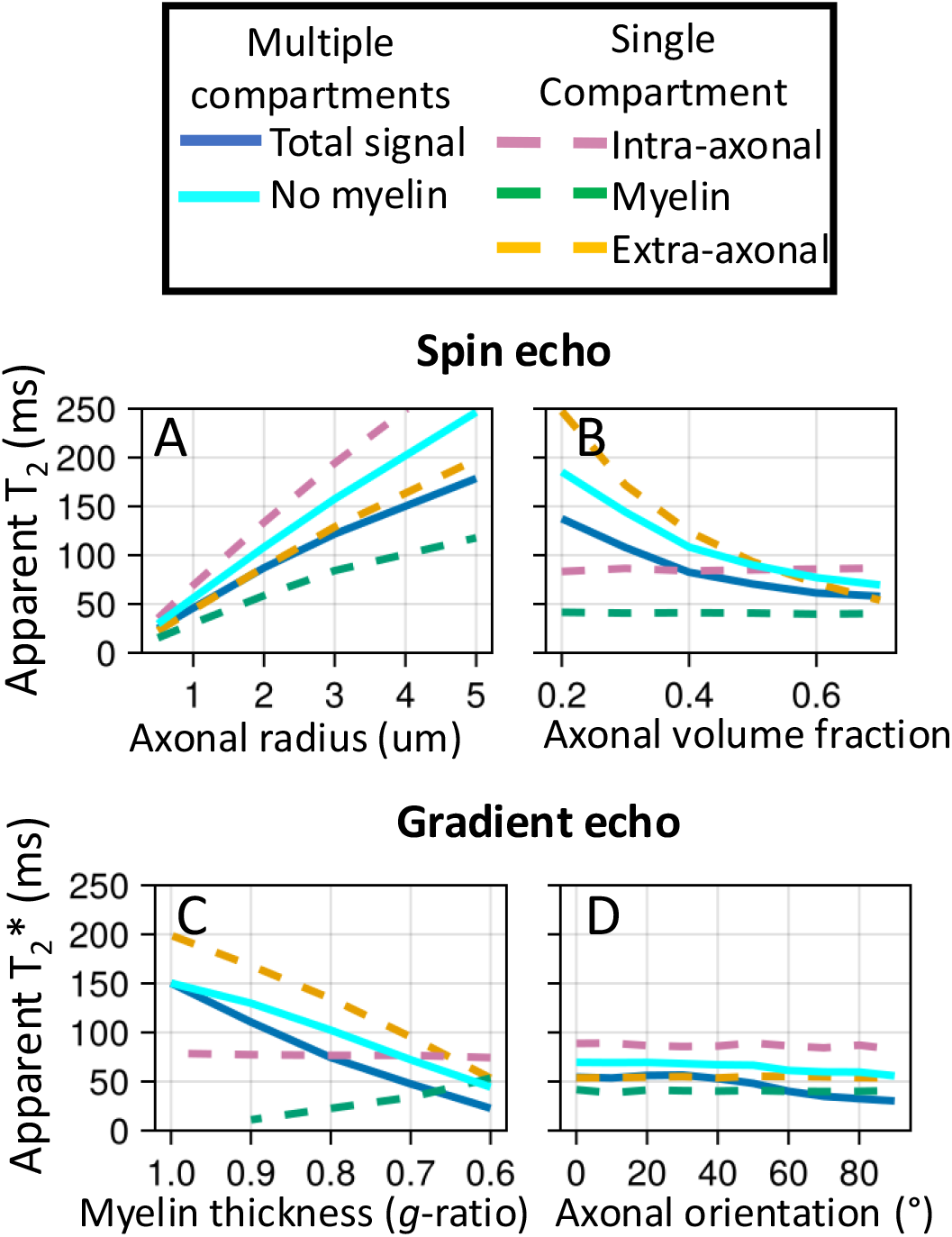
The trend of T2 in a spin echo experiment (top) and T2* in a gradient echo experiment (bottom) as we change different aspects of the microstructure. The baseline microstructure and sequences are the same as in **Figure 8**. The simulations include both the surface relaxation and the myelin-induced off-resonance field, of which the effects are illustrated in Figure 8. A) Effect of increasing axonal radius (when all axons have the same radius); B) effect of changing the axonal volume fraction (while maintaining the axon radius); C) the effect of myelin loss from the outside in (i.e., myelin is replaced by extra-axonal space); D) effect of the angle between the main magnetic field and the axon orientation.

Increasing the myelin thickness increases the frequency offset between compartments and lowers the T_2_* (Figure 9C) as previously noted by Xu *et al*. (2018). Note that the dependence of the T_2_* of the intra-axonal or myelin water on the myelin thickness is not caused by the myelin-induced off-resonance field, but is rather caused by the space for the myelin water increasing at the expense of the extra-axonal space, which will affect the T_2_ due to surface relaxation.

Finally, we illustrate how the T_2_* depends on the angle between the axons and the main magnetic field (Figure 9D). This matches previous simulations (Chen, Foxley and Miller, 2013; Kor *et al*., 2019) and has also been observed in the brain’s white matter (Bender and Klose, 2010; Denk *et al*., 2011; Lee *et al*., 2011).

The simulator enables the analysis of how microstructural changes affect the MRI signal across multiple modalities. To illustrate this Figure 10 shows how the signal changes across the four modalities discussed in this section (diffusion-weighted MRI, magnetisation transfer, spin echo, and gradient echo). The first two panels consider halving the myelin density, either by halving the fraction of myelinated axons (from 0.5 to 0.25) or by halving the thickness of the myelin sheaths (increasing the *g*-ratio from 0.7 to 0.81). Both have a very similar profile of signal change across all the considered modalities, which closely matches the profile seen for axonal loss (central panel). On the other hand, doubling the axonal diameter does have a different profile as it affects the spin echo, gradient echo, and MTR signal, but not the diffusion-weighted MRI (at the *b*-value considered here). Changing the axon orientation with respect to the main magnetic field changes the gradient echo (due to a changing off-resonance field generated by the myelin), but not the spin echo or MTR.

**Figure 10.**
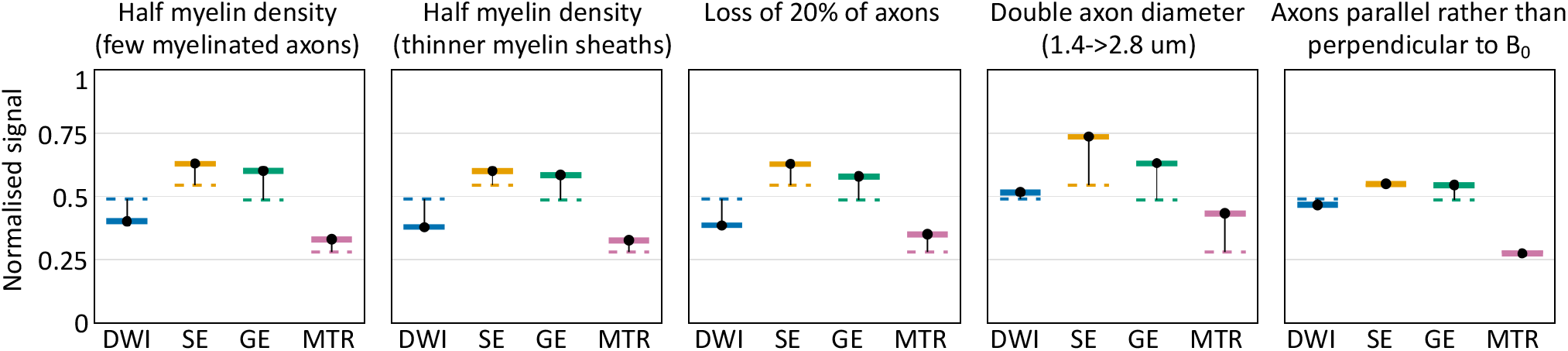
How microstructural changes affect the signal for 4 MRI modalities: diffusion-weighted imaging (DWI), spin echo (SE), gradient echo (GE), and magnetisation transfer ratio (MTR). These are the same sequences that were used in Figure 5 for DWI, Figure 7 for MTR, and Figures 8 and 9 for SE and GE. The DWI uses a b-value of 1 ms^2^/um with gradients orthogonal to the axons. The SE and GE both have an echo time of 50 ms. For simplicity, instantaneous excitation and refocus RF pulses were used for all sequences, however the saturation RF pulses used in MTR are finite. The baseline signal (dashed lines) was based on randomly packed parallel axons with an inner diameter of 1.4 um, half of which are myelinated with a g-ratio of 0.7 (leading to an outer diameter of 2 um). Each panel shows how this baseline signal changes for each sequence in a specific scenario.

## 4. Discussion

We present a new MRI Monte Carlo simulator that captures the effect of T_1_/T_2_ relaxation, diffusion, permeability, off-resonance fields, surface relaxivity, and magnetisation transfer on a given MRI sequence. The user provides a representation of the tissue of interest, biophysical parameters of interest (Table 1), and the MRI sequence(s) to be simulated.

MRI sequences can be produced using the accompanying MRIBuilder.jl Julia library or any of the existing tools for generating pulseq files (Layton *et al*., 2017), such as the python pypulseq or the matlab pulseq library. Realistic tissue geometries beyond cylinders or spheres are more difficult to obtain. We do not, at present, provide a tool for this, but note that there are several computational tissue phantom generators available (Ginsburger *et al*., 2019; Palombo, Alexander and Zhang, 2019; Callaghan *et al*., 2020; Villarreal-Haro *et al*., 2023). Meshes produced by any of these tissue generators can be used as inputs to MCMR simulator.

By varying the parameters in these tissue generators one can use the simulator to design sequences sensitive to specific aspects of the tissue microstructure as well as to train a model to fit any data generated from these sequences. The multi-modal nature of the simulator would allow an AI model generated in this way to combine information across multiple MRI modalities. We plan to explore such applications in future work.

In the future such computational tissue phantom generators would ideally not just focus on creating a close approximation of healthy tissue, but also simulate how this tissue might change due to, for example, inflammation or cell death. The simulator could then be used to predict how these would affect the signal across a wide range of MRI modalities, which could then be compared with the pattern of change observed in actual tissue in different disease states (Rafipoor *et al*., 2022).

Running the simulations for the results took about two hours on a standard laptop. Larger simulations can be trivially parallelised across multiple cores or multiple jobs submitted to a computing cluster. The simulator runtime is given by the number of spins multiplied by the number of timesteps multiplied with the duration of each timestep for each spin. The duration of each timestep will be dominated by the time needed to compute the off-resonance field, if present, or the collision detection otherwise. One exception is when running more than ∼100 sequences simultaneously, when the magnetisation update for all of those sequences might become rate-limiting.

The main ways for a user to speed up the simulator are to reduce the number of timesteps or the number of spins. Both of these have a different effect on the predicted signal. A sufficiently small timestep will have no effect on the signal, but if it is too large it might cause a systematic bias. While the simulator includes multiple heuristics to select an optimal timestep (S1), the resulting timestep might be too conservative for many MRI sequences.

For example, by default the timestep is kept short enough that one can accurately estimate the tortuosity in the extra-cellular space, however if the sequence in question is not sensitive to this tortuosity, then the timestep could be greatly increased without biasing the results. When running a small number of simulations, we recommend the use of the default, conservative timestep. When running many simulations, we recommend to vary the timestep in a test case to find out above which timestep there is a systematic bias in the predicted signal. Reducing the timestep below this value will not give additional benefits.

The effect of the number of spins on the total signal is different to that of the timestep. Decreasing the number of spins does not cause a systematic bias (i.e., does not affect the accuracy), but can increase the amount of random noise (i.e., decreases the precision). The size of this random noise is hard to predict beforehand. It depends on how many spins actually contribute to the final signal, which might actually be a small fraction of the total number of spins. As this is in essence a counting experiment (where we count the number of contributing spins), we can estimate the signal-to-noise ratio to be roughly 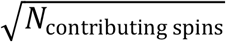. If the attenuation is small, we can be sure that the number of contributing spins is large. However, if the attenuation is large, the opposite is not necessarily true. For example, in a spin echo experiment the attenuation might be small because there are just a few spins with a long T_2_ (i.e., few contributing spins) or because all the spins have the same short T_2_ (i.e., a lot of spins equally contributing a small amount). Accurately, simulating the former case requires many more spins than the latter to ensure that there are enough with a long T_2_. As a practical strategy, one can estimate the noise level by running the simulation a few times with a small number of spins and different initial seeds. One can then estimate how many spins would need to be simulated to achieve a sufficiently low noise level for the desired application using the fact that the noise level decreases with the square root of the number of spins.

The simulator effectively captures effects on length scales on the order of 1-100 μm and timescales on the order of milliseconds to seconds. Smaller-scale effects (tumbling, spin-spin interactions, spin-lattice interactions) can be effectively included by setting T_1_ and T_2_. The simulator could in the future be extended to produce an MR image covering the full field of view rather than just a single voxel. Such an extension is made possible by the capability of the simulator to predict the MRI signal for multiple sequences at once. Instead of these actually representing different sequences, the different sequences could be used to represent how the same sequence would look like for different locations in the brain by varying the origin of the magnetic field gradients and the strength of the RF pulse.

Possible future extensions to increase the realism of the MRI signal predictions include:

- Varying proton density between different compartments.
- Varying diffusivities between different compartments (i.e., tortuosity).
- More realistic magnetisation transfer by allowing more than a single mobile and single bound pool. This would allow for the modelling of a mobile macromolecular pool as well as the inclusion of multiple surface-bound pools (e.g., separately representing water bound by surface tension as well as the magnetisation of membrane macromolecules; Manning, MacKay and Michal, 2021).
- Adding time-dependent transverse relaxation, which would more accurately capture the spectral lineshape of macromolecules (Assländer *et al*., 2022; Sukstanskii and Yablonskiy, 2023).
- Allowing the T_2_ of the surface-bound pool to depend on membrane orientation with respect to the main magnetic field to correctly model orientation-dependent magnetisation transfer (Pampel *et al*., 2015).
- Addition of specific metabolites or peptides, which are detected in MR spectroscopy or targeted in Chemical Exchange Saturation Transfer (CEST).
- Allowing tissue to vary during the MRI acquisition. This could be used to vary the geometry (e.g., to model motion or cell swelling) or the magnetic susceptibility (e.g., to model variations in blood oxygenation).

The simulator was developed using modern software practices including thousands of unit and integration tests. Extensive documentation including installation instructions and a tutorial is available at https://michielcottaar.github.io/MCMRSimulator.jl/stable/.

## Acknowledgements

M.C., S.J. and K.L.M. are supported by the Wellcome Trust (203139/Z/16/Z, 215573/Z/19/Z, 221933/Z/20/Z, 224573/Z/21/Z). BCT is funded by a Sir Henry Wellcome Postdoctoral Fellowship (Wellcome Trust; 222829/Z/21/Z). Z.Z. is supported by the University of Oxford and China Scholarship Council. This project was supported by the NIHR Oxford Health Biomedical Research Centre (NIHR203316). The views expressed are those of the author(s) and not necessarily those of the NIHR or the Department of Health and Social Care. The Wellcome Centre for Integrative Neuroimaging is supported by core funding from the Wellcome Trust (203139/Z/16/Z and 203139/A/16/Z). For the purpose of open access, the author has applied a CC BY public copyright licence to any Author Accepted Manuscript version arising from this submission.

## Supplementary Information

### S1. Determining the simulations timesteps

To accurately handle all of the features discussed in the main paper’s theory section there are many constraints on the timestep of the simulator. These constraints include those imposed by the sequences, by the geometry, and by having a sufficient rate of collisions (i.e., short enough timestep) to accurately model permeability, surface relaxation, and magnetisation transfer.

If the timestep is too long for any of these constraints this could bias the final results. All of these constraints could be satisfied simply by using a very short timestep, which would be very computationally expensive. Here we discuss the algorithm used in the simulator to select the maximum timestep that can be used to obtain accurate simulation results with minimal computational costs.

Firstly, we split the total sequence (or multiple sequences) based on the start or end of any RF pulses or building blocks used in the sequence definition, as well as the timings of any instantaneous RF pulses, gradients, or readouts. At this stage we also ensure that within each period the gradient can be approximately described as changing linearly. This ensures that during any timestep the sequence is relatively homogeneous (e.g., there is a pulse active or there isn’t) and that all of the instantaneous events are at the edges of any timestep.

Secondly, we iterate through each of the time periods from the split defined above and for each determine whether it should be split up any further. This is done by determining at each time period the maximum timestep τ_ma*x*_. The total time period is then split up in uniform steps, where each step duration is shorter than τ_ma*x*_.

The maximum timestep τ_ma*x*_ is set to the minimum of various terms. The constants in front of each term have been empirically determined as illustrated in Figure S1 and Figure S2:

1. 0.03 *l*^2^/*D*, where *l* is the minimum size scale of the obstructions included in the tissue microstructure phantom and *D* is the diffusivity. This reflects that for timesteps larger than this the ability of spins to travel around these obstructions is reduced (i.e., the tortuosity becomes timestep-dependent). The value of 0.03 is set based on simulations of this tortuosity (Figure S1). It also ensures that within compartments it is not consistently the same spin that undergoes all the collisions, but that instead the collision rate is equally shared across all spins (not shown). For cylinders and spheres the size scale is set to the radius, and for walls, to the distance between neighbouring walls. For meshes it is set by estimating the equivalent radius based on the local curvature of the mesh.
2. 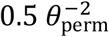, which ensures that the permeability is accurately modelled (Figure S2B).
3. 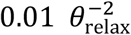, which ensures that the surface relaxation is accurately modelled (Figure S2C).
4. 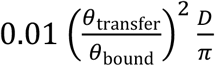, which ensures that the transition from free to bound states is accurately modelled (Figure S2D).
5. 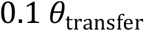, which ensures that the same spins that have just been released from the bound state can return in a time short compared with the bound spin dwell time (*θ*_transfer_) to get bound again at the same surface.
6. 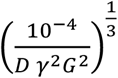, where γ is the gyromagnetic ratio and *G* is the sequence gradient strength. This ensures that a spin taking a typical step size 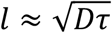 given the diffusivity and timestep will not move so much that the phase accumulation during that timestep would be very different at the beginning or end of the timestep (i.e., γ*Gl*τ ≪ 1).

Note that the final constraint depends on the sequence, so the maximum timestep can be different at different times during the sequence evolution.

**Figure S 1.**
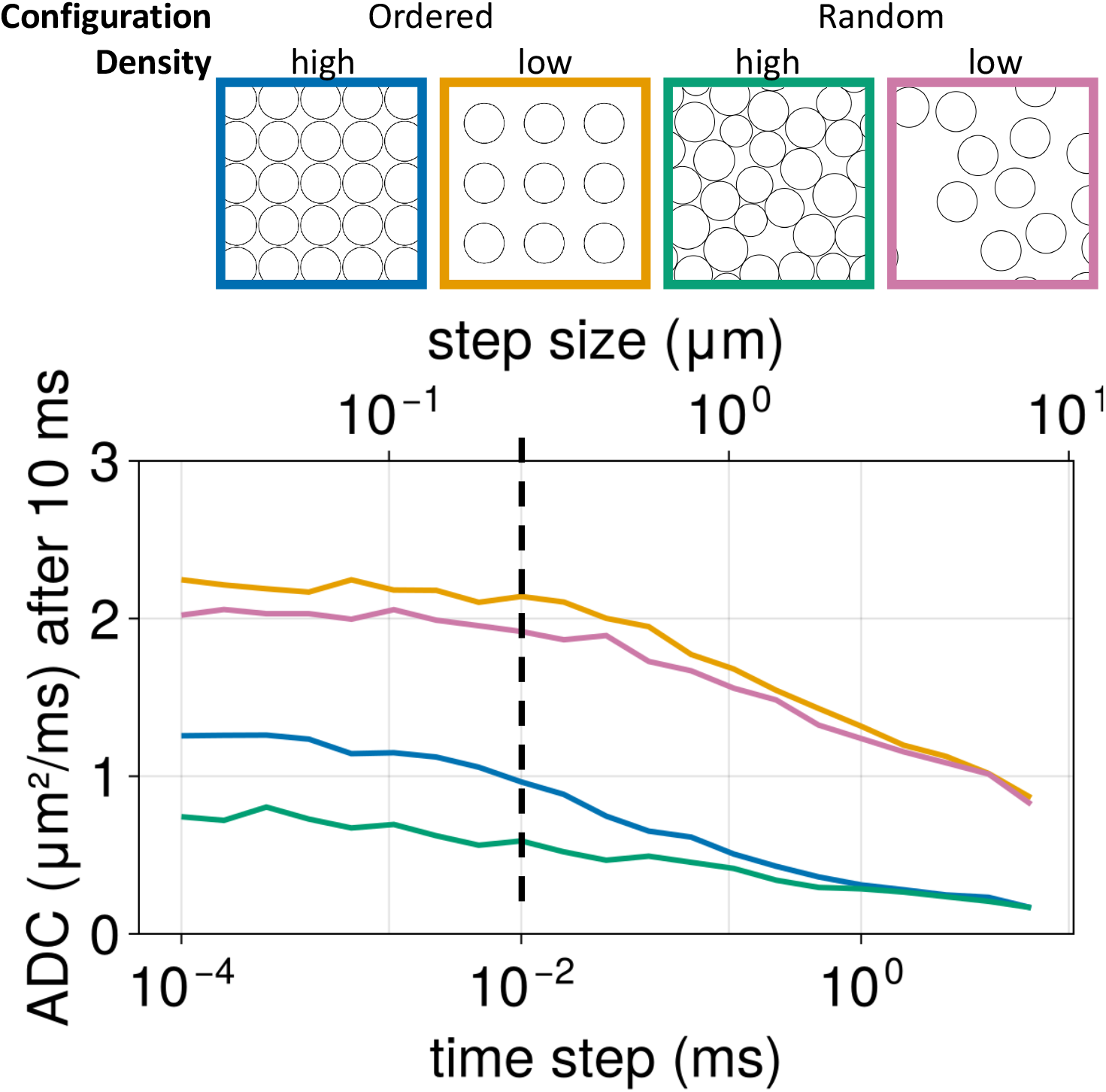
Constraint on the timestep of the simulator in order to accurately model the tortuosity in the extra-cellular space. We test this case by considering various cylinder (r = 1μm) packing configurations with different levels of orderedness and density (top). The intrinsic diffusivity is 3 μm/ms^2^, however this diffusivity is reduced orthogonal to the axons to some apparent diffusion coeffecient (ADC) due to the tortuosity of spins trajectories being hindered by the cylinders. For all simulated packing configurations we find that the diffusivity of spins orthogonal to these cylinders becomes timestep-independent for timesteps shorter than 0.01 ms. This can be extended to different radii and intrinsic diffusivities as 0.03 l^2^/D, where l is the cylinder radius and D is the diffusivity.

Figures S2 compares three different methods to make the permeability, surface relaxation, and magnetisation transfer timestep-independent. For all metrics these three methods converge for small timesteps, however we want to adopt the method that retains same result the longest as the timestep increases. For surface relaxation and magnetisation transfer this is the logarithmic adjustment (orange). For permeability this is also a logarithmic adjustment, but with the addition of a Bessel function (green). The result of these simulations is also used to set the maximum timestep allowed to accurately model these effects as listed above.

**Figure S 2.**
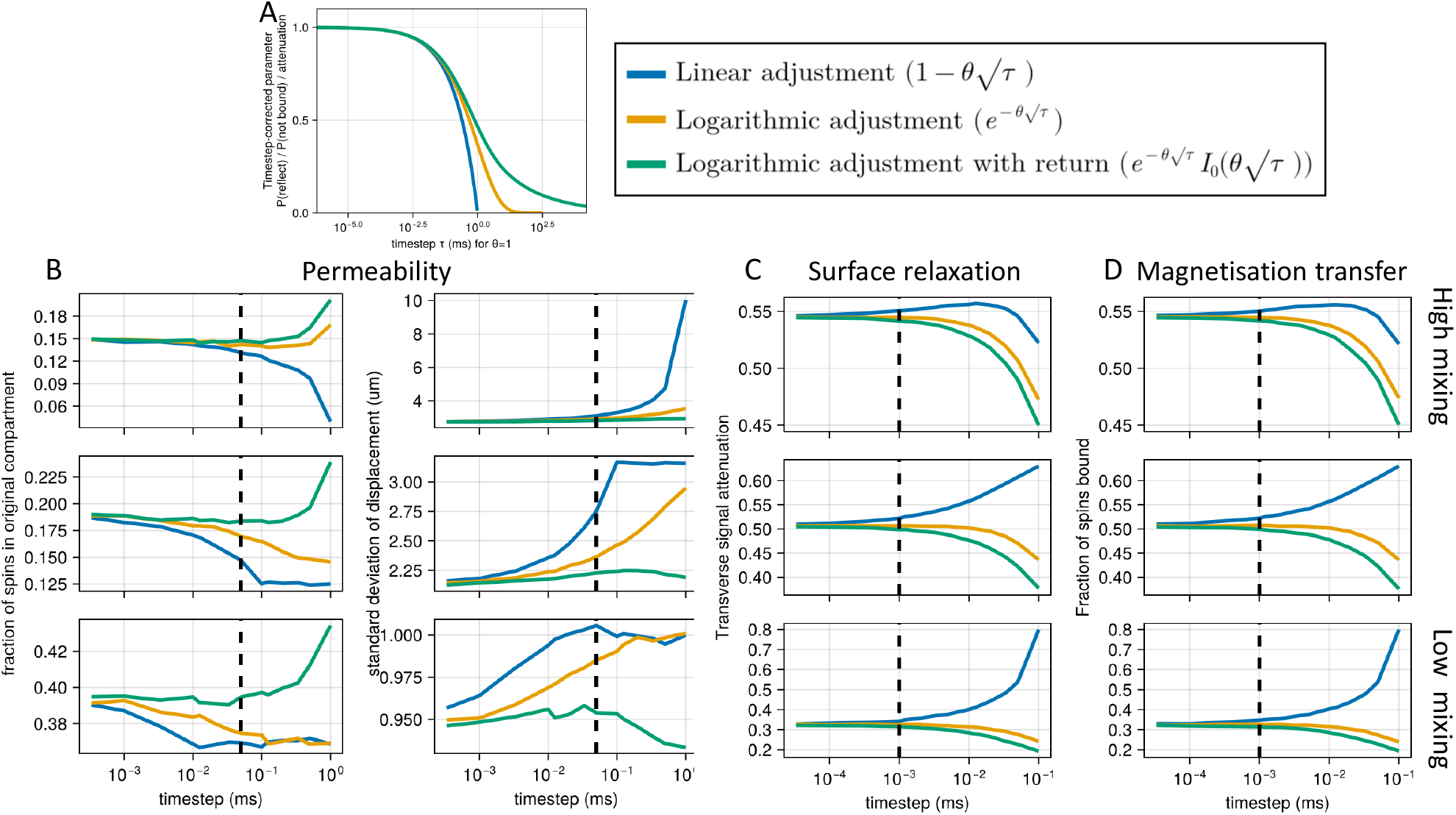
Three different methods to correct for the timestep-dependence of the number of collisions (A) and how they affect the permeability (B), surface relaxation (C), and magnetisation transfer (D) in a simple toy example. This toy example considers 1-dimensional diffusion with obstructions regularly spaced every 1 μm. These obstructions may be permeable (B), have surface relaxation (C), or magnetisation transfer to a bound pool (D). For permeability we compare 2 metrics (fraction of spins in origin compartment and standard deviation of the displacement). For the other parameters we only consider a single metric (signal attenuation for surface relaxation and fraction of spins transferred from free to bound pool for magnetisation transfer). In B-D the top row shows an example for well-mixed compartments, while at the bottom the exchange/relaxation/transfer rates are short compared with the mixing times. The first correction method (blue) is derived from the naïve assumption that as the rate of collision doubles due to timestep changes, the effect of each collision should be halved. The second (orange) makes the same assumption, but halves the effect size in logarithmic space rather than linear space, which is appropriate for the surface relaxation and magnetisation transfer (C, D). Finally, the third (green) adds the Bessel function to take into account that two instances of the spin passing through a permeable membrane will effectively cancel each other out (B). The black dashed lines show the maximum timestep adopted by default in the simulator.

### S2. Timestep-dependent collision rate

Here we derive the collision rate of spins with the surface in the simulations. This collision rate is crucial to correctly derive the exchange rate, surface relaxation, and magnetisation transfer rate within the simulator.

We will first derive this collision rate for the case of a single infinite wall before arguing why this result is more generally valid. In the case of the infinite wall, we only have to consider the position along a single dimension, namely along the wall normal. We refer to this dimension as *x*. The probability of a spin being between positions *x* and *x* + *dx* is given by the line density *λ* multiplied with *dx*.

The probability of any random spin at distance *x* hitting the wall in the next timestep (with step size 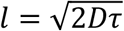) is given by:

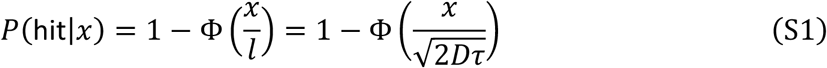

Where Φ is the cumulative distribution of the normal function:

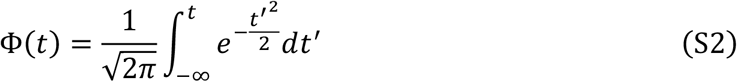

Using this we can compute the total number of spins hitting the wall from one side as

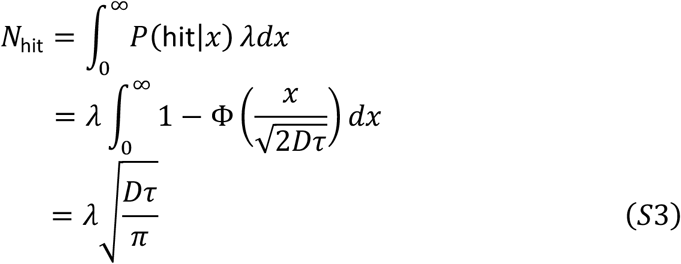

We can convert this to a finite patch of the wall with surface area *S* by using *λ* = *ρS*, where *ρ* is the volumetric density:

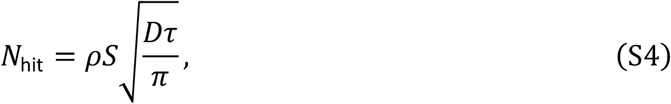

Finally, we consider that the spins are within a large voxel with volume *V* and total number of spins of *N*_spins_:

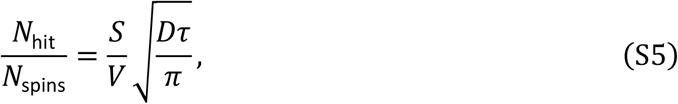

If this is the number of spins hitting the wall during each timestep τ, the collision rate *r*_*c*_ is given by:

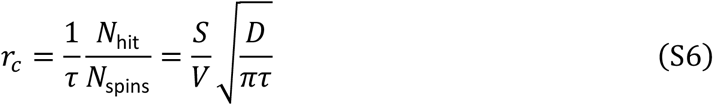

The above derivation assumes that spins can hit the wall from infinitely far (as we integrate to infinity). If there are other obstructions in the way then this will not hold. However, the number of spins prevented from hitting the wall by other obstructions, will be exactly replaced by the number of spins hitting the wall after bouncing of these other obstructions. So, the total rate of collisions is unaffected by the presence of other obstructions. A similar logic can be used to conclude that both permeability and magnetisation transfer also do not affect the collision rate.

